# Meiosis Initiates In The Fetal Ovary Of Mice Lacking All Retinoic Acid Receptor Isotypes

**DOI:** 10.1101/716498

**Authors:** Nadège Vernet, Manuel Mark, Diana Condrea, Betty Féret, Muriel Klopfenstein, Violaine Alunni, Marius Teletin, Norbert B. Ghyselinck

**Affiliations:** Institut de Génétique et de Biologie Moléculaire et Cellulaire (IGBMC), Département de Génétique Fonctionnelle et Cancer, Centre National de la Recherche Scientifique (CNRS UMR7104), Institut National de la Santé et de la Recherche Médicale (INSERM U1258), Université de Strasbourg (UNISTRA), 1 rue Laurent Fries, BP-10142, F-67404 Illkirch Cedex, France; Service de Biologie de la Reproduction, Hôpitaux Universitaires de Strasbourg (HUS), France; GenomEast platform, France Génomique consortium, IGBMC, 1 rue Laurent Fries, F-67404 Illkirch Cedex, France

## Abstract

Gametes are generated through a specialized cell differentiation process, meiosis which, in most mammals, is initiated in ovaries during fetal life. It is widely admitted that all-*trans* retinoic acid (ATRA) is the molecular signal triggering meiosis initiation in mouse female germ cells, but a genetic approach in which ATRA synthesis is impaired disputes this proposal. In the present study, we investigated the contribution of endogenous ATRA to meiosis by analyzing fetuses lacking all RARs ubiquitously, obtained through a tamoxifen-inducible cre recombinase-mediated gene targeting approach. Efficient ablation of RAR-coding genes was assessed by the multiple congenital abnormalities displayed by the mutant fetuses. Unexpectedly, their germ cells robustly expressed STRA8, REC8, SYCP1 and SYCP3, showing that RAR are actually dispensable up to the zygotene stage of meiotic prophase I. Thus our study goes against the current model according to which meiosis is triggered by endogenous ATRA in the developing ovary and revives the identification of the meiosis-preventing substance synthesized by CYP26B1 in the fetal testis.

## Introduction

Mammalian meiosis is a germ-cell specific division process that generates haploid gametes from their diploid precursors, oogonia in the female and spermatogonia in the male. In the mouse, female germ cells enter into meiosis before birth, around embryonic day 13.5 (E13.5)^1^. During the same embryonic period, male germ cells stop proliferating, and enter the G0/G1 phase of the cell cycle thus becoming mitotically quiescent. Male germ cells resume proliferation at birth, and then enter into meiosis starting from post-natal day 8^2^.

In order to account for the sexual dimorphism in the timing of gem cell differentiation, it was hypothesized, notably from transplantation experiments of germ cells^3^, that the decision to enter meiosis is controlled by a meiosis-inducing substance (MIS) or/and by a meiosis-preventing substance (MPS) produced by somatic cells^4^. Subsequently, it was proposed that all-trans retinoic acid (ATRA) and its degrading enzyme CYP26B1 played key roles in controlling the timing of meiosis initiation in female and male gonads, respectively^5,6^. The concept that ATRA is the MIS was however challenged by a genetic study demonstrating that meiosis initiation occurs despite the lack of two major ATRA-synthesizing enzymes^7^. The recent finding that a third enzyme, expressed in fetal ovaries and capable of ATRA synthesis, is also involved in meiosis revived the model according to which meiosis entry is triggered by endogenous ATRA in the ovary^8^

ATRA is the active metabolite of retinol (vitamin A). Inside cells, conversion of retinol to ATRA depends upon retinaldehyde dehydrogenases (ALDH1A1, ALDH1A2 and ALDH1A3 isotypes)^9^. Then, ATRA activity is mediated by the nuclear retinoic acid receptors (RARA, RARB and RARG isotypes), which are ligand-dependent transcriptional regulators. They usually function in the form of heterodimers with rexinoid receptors (RXRs) to control expression of ATRA-target genes, in which they are bound to specific DNA sites called retinoic acid response elements (RARE)^10^. In addition, RAR are capable of non-genomic activation events at the cell membrane^11^, similarly to steroid nuclear receptors^12^.

As all ATRA-dependent events rely in some way on RAR, we decided to tackle the potential contribution of endogenous ATRA to meiosis in female germ cells by generating and analyzing mice lacking all RAR-isotypes. Against the currently admitted model^13,14^, our study reveals that RARs (and therefore endogenous ATRA) are in fact fully dispensable for meiotic initiation in the mouse fetal ovary.

## Results

### Expression of RARs in the fetal ovary

The expression of *Rars* in the fetal gonads is poorly documented^15–17^. To determine which RAR isotypes are actually present in the ovary, we performed immunohistochemistry (IHC). At E11.5, RARA was detected in a large number of tissues, including the somatic cells of the ovary but not germ cells (Fig. 1A,C-E). No information was obtained for RARB since reliable antibodies for RARB are not available^18^. RARG was readily detected in cartilages, but not in the ovary (Fig. 1B-B’). Then we performed RT-qPCR on single ovarian cells at E13.5 (n=25) and E14.5 (n=40), to which the germ cell identity was assigned based on the expression of *Dazl*, *Ddx4* and *Kit*. *Rara* and *Rarg* mRNAs were detected in a majority of germ cells at E13.5 and E14.5 (Fig. 1F,G). No information was obtained for *Rarb* mRNA since the mice we used were on a *Rarb*-null genetic background (see Methods). It is therefore evident that at least two RAR mRNAs are present in female germ cells, but at some stage their level of expression is below the threshold of detection by IHC.

**Fig. 1.**
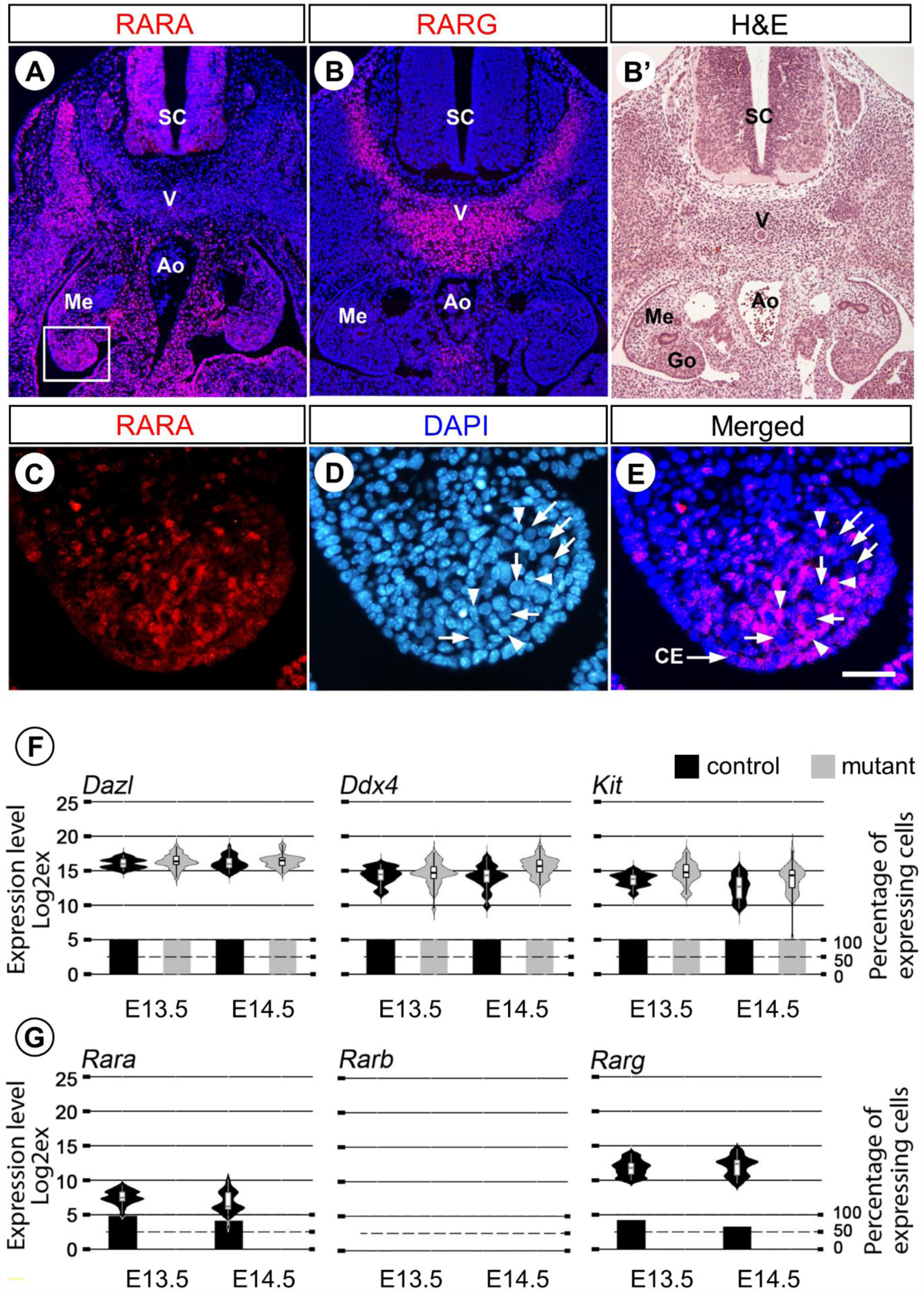
Expression of RARs in the female gonad and germ cells during embryonic development. (A and B) Immunohistochemical detection of RARA and RARG (red signals) on frontal histological sections of a E11.5 wild-type female embryo: expression of RARG is confined to the precartilaginous anlage of a vertebra, while that of RARA is more widespread and includes notably the undifferentiated gonad. (B’) Same section as (B) stained with hematoxylin and eosin. (C-E) Enlargement of the box in (A): RARA (in red) is detected in the nuclei of some somatic cells of the gonad (arrowheads) and of the coelomic epithelium; in contrast, the large, rounded nuclei characteristic of germ cells (arrows) do not exhibit anti-RARA immunostaining. Nuclei are counterstained with DAPI (blue signal). Ao; aorta; CE, coelomic epithelium; Go, gonad; Me, mesonephros; SC, spinal cord; V, vertebra; Scale bar (in E): 160 µm (A,B,B’) and 30 µm (C-E). (F,G) RT-qPCR analysis comparing the expression levels and distributions of the germ cell markers *Dazl*, *Ddx4* and *Kit* (F) and *Rara*, *Rarb* and *Rarg* (G) mRNAs in single germ cells from control and mutant ovaries at E13.5 and E14.5. The Violin plot width and length represent respectively the number of cells and the range of expression (Log2Ex). The box-and-whisker plots illustrate medians, ranges and variabilities of the collected data. The histograms show the percentages of expressing cells in each group. At respectively E13.5 and E14.5, *Rara* is present in 96% and 83% of germ cells; *Rarb* in none of germ cells since the fetus are on a *Rarb*-null genetic background (see Methods); *Rarg* is present in 84% and 65% of germ cells. Importantly, *Rara*, *Rarb* and *Rarg* are undetectable in germ cells isolated from the *Rar*-mutant ovaries (see main text for details).

### Efficient ablation of all RARs in the developing gonad from E11.5 onwards

Given the expression pattern of RARs, we reasoned that full impairment of ATRA signaling in the whole fetal ovary would require the ablation of all three RAR-coding genes. This was not possible by associating *Rara*, *Rarb* and *Rarg* knockout alleles in a single fetus, because *Rara*^−/−^;*Rarg*^−/−^;*Rarb*^+/−^ embryos do not develop beyond E8.5, precluding analysis of their ovaries^19^. To do so, we performed cre-directed genetic ablation of *Rara* and *Rarg* using a ubiquitously expressed cre/ERT^2^ activated by TAM, prior to meiotic initiation, but later than E8.5, in the context of a *Rarb*-null background (see Methods). We first chose to administer TAM at E10.5, shortly after the start of gonad formation, but three days before meiosis initiation. Ablation of RARA and RARG was assessed by IHC at E11.5, i.e., 24 hours after TAM-induction of cre/ERT^2^. Immunostaining for RARG was nearly abolished in mutant embryos in all RARG-expressing tissues. Only a few cells reacted with the antibody (Suppl. Fig. 1). At E14.5, the number of cells remaining positive for RARG in mutants was even smaller, while no cell expressed RARA (Suppl. Fig. 2). Thus efficient ablation of RARA and RARG was widespread at E11.5 and complete at E14.5.

The pattern of gene excision by cre/ERT^2^ was assessed using the *mT/mG* reporter transgene in control and mutant fetuses (see Methods). Expression of mGFP was detected in all germ cells of the mutants at E11.5, and in almost all of them at E14.5 (Suppl. Fig. 3). This indicated that cre/ERT^2^-directed excision of the reporter occurred in almost all germ cells, as early as 24 hours after TAM treatment. In agreement with this finding, *Rara* and *Rarg* mRNAs were not detected in any germ cell isolated from mutant ovaries at E13.5 or E14.5, and analyzed by RT-qPCR (Fig. 1G). Altogether, these data showed that efficient excision of all RARs occurred in all germ cells.

### Ablation of all RARs does not impair meiosis initiation

To investigate the impact of RAR ablation on meiotic initiation, the expression of canonical markers of meiosis was assessed by IHC throughout the anteroposterior axis of the ovary. At E14.5, numerous germ cells in both control and mutant fetuses expressed the synaptonemal protein SYCP3^20^ and the cohesin REC8^21^ (Fig. 2A). The mean number of germ cells, and the percentages of SYCP3-positive or REC8-positive germ cells were similar in control and mutant fetuses (Fig. 2B,C). This indicated that meiosis initiated in germ cells of the mutant fetuses.

**Fig. 2.**
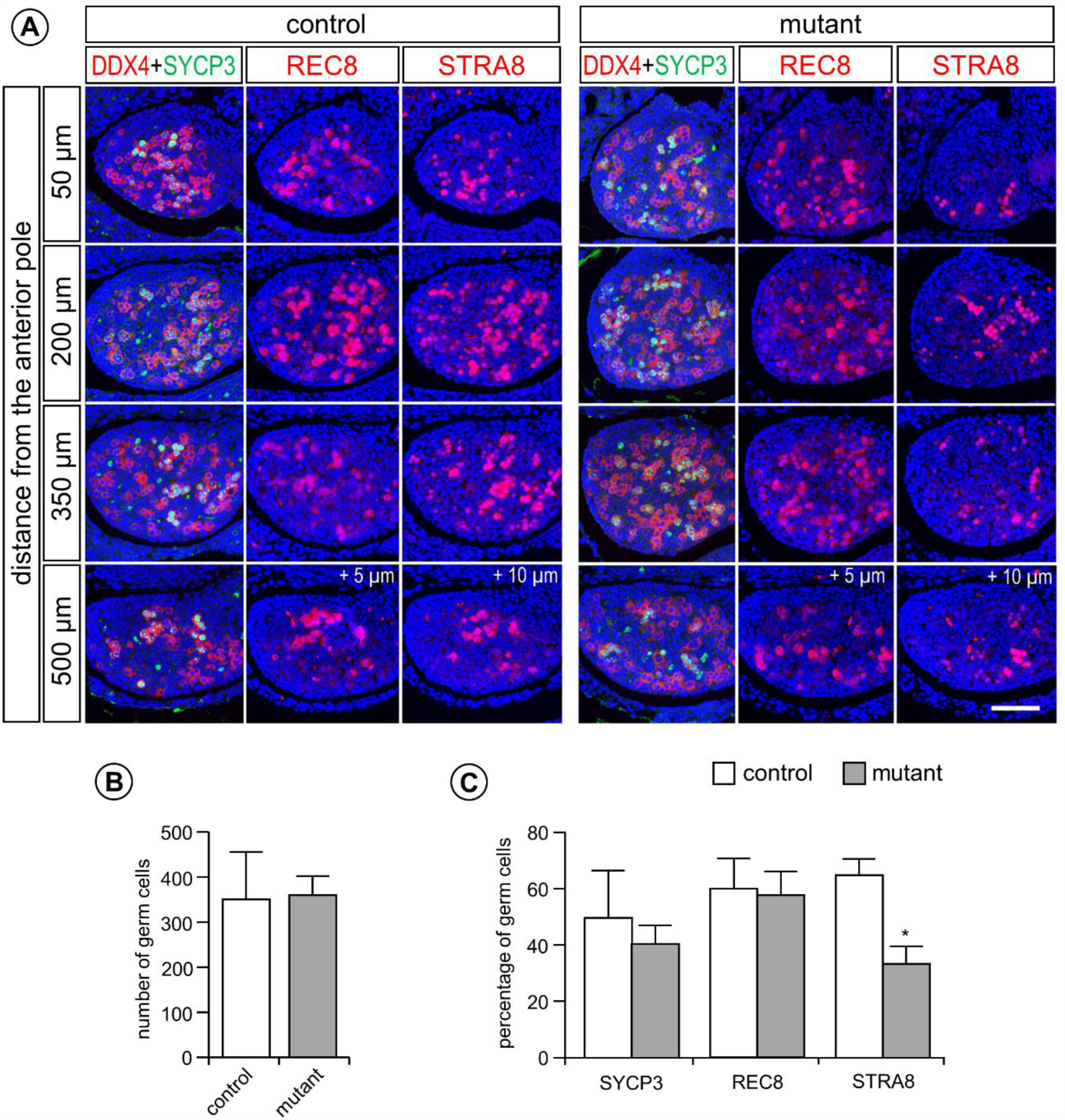
Markers of meiotic prophase I are robustly expressed at E14.5 in ovaries of mutants lacking RARs. (A) Detection of meiotic cells expressing SYCP3 (green nuclear signal), REC8 or STRA8 (red nuclear signals) on consecutive, 5 µm thick, transverse histological sections at four different levels of the ovaries from control and mutant fetuses, as indicated. DDX4 (red cytoplasmic signal) is present in all germ cells. The positions of histological sections along the anteroposterior axis is indicated in terms of distance from the anterior pole of the ovary (i.e., 50, 200, 350 and 500 microns). + 5 µm and + 10 µm: the indicated histological sections “REC8” and “STRA8” on each line are, respectively, 5 and 10 microns apart from the indicated section “DDX4 + SYCP3” that was used to establish the total number of germ cells. Nuclei are counterstained with DAPI (blue signal). Scale bar: 60 µm. (B) Average of the total number of germ cells present at the 4 different levels of the ovary illustrated in panel A in 4 control (white bars) and 4 mutant (grey bars) fetuses at E14.5. (C) Mean percentages of germ cells expressing SYCP3, REC8 and STRA8 in 4 control (white bars) and 4 mutant (grey bars) fetuses at E14.5. The asterisk (in C) indicates a significant difference (p<0.05).

To rule out the possibility that the cells which initiated meiosis experienced an inefficient ablation of the RAR-coding genes, we took advantage of the presence of *mT/mG* reporter in the fetuses. The vast majority of the SYCP3- or REC8-positive germ cells of the mutant ovary were mGFP-positive (Fig. 3A-F), indicating excision in almost all of the meiotic cells. Surprisingly, mGFP was never detected in somatic cells of the ovary nor in the mesonephros, making questionable whether cre/ERT^2^ was efficient or whether the reporter was expressed in these cells. To assess RAR ablation in somatic cells and exclude the possibility that cre-mediated excision was mosaic, we performed genomic PCR analysis, using DNA extracted from whole ovaries of E13.5 control and mutant fetuses. Excised (L-), but not conditional (L2), alleles of *Rara* and *Rarg* were detected in genomic DNA isolated from mutant ovaries. In contrast, conditional (L2), but not excised (L-), alleles were always detected in control ovaries (Fig. 3G,H). Altogether, these data show that efficient excision of all RARs occurs in all of their tissues, including somatic and germ cells in the ovaries.

**Fig. 3.**
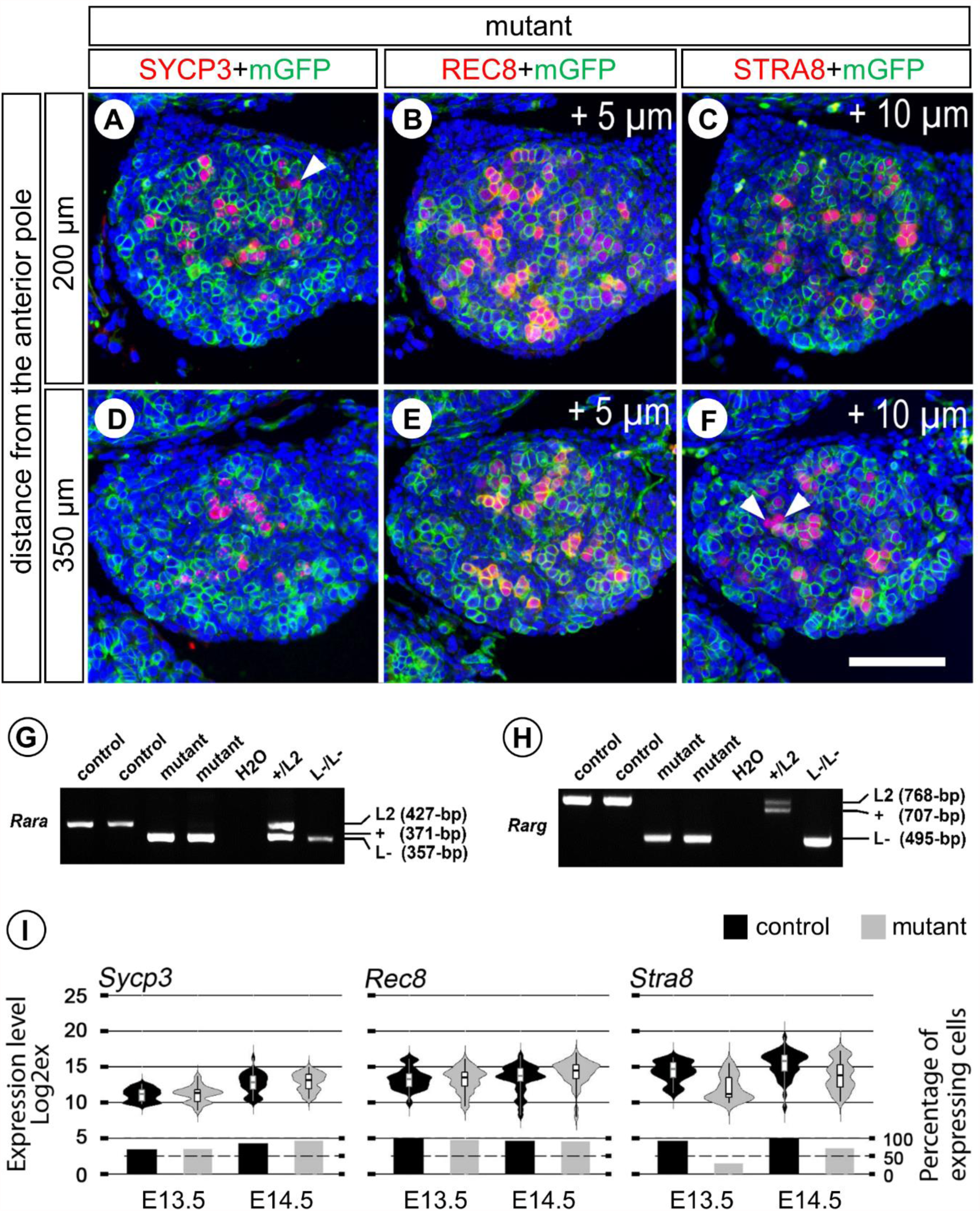
Evidence that gene excision has actually occurred in meiotic cells 4 days after administration of TAM. (A-F) Detection of the meiotic markers SYCP3, REC8 or STRA8 (red nuclear signals) and of mGFP (green membranous signal) on consecutive, 5 µm thick, transverse histological sections at two different levels of the ovary of a mutant fetus at E14.5. Efficient excision of the reporter transgene by cre/ERT^2^ is assessed by mGFP expression in virtually all meiotic germ cells. Possible exceptions (i.e, red nuclei without a green contour) are indicated by white arrowheads. The position of histological sections along the anteroposterior axis is indicated on the left side in terms of distance from the anterior pole of the ovary (i.e., 200 and 350 microns). + 5 µm and + 10 µm: the indicated histological sections “REC8 + mGFP” and “STRA8 + mGFP” on each line are, respectively, 5 and 10 microns apart from the indicated section “SYCP3 + mGFP”. Nuclei are counterstained with DAPI (blue signal). Scale bar (F): 60 µm. (G,H) PCR analysis of genomic DNA extracted from ovaries of control and mutant fetuses at E13.5, as indicated. (G) DNA was genotyped using primers 5’-CAGGGAGGATGCTGTTTGTA-3’, 5’-AACTGCTGCTCTGGGTCTC G-3’ and 5’-TACACTAACTACCCTTGACC-3’ to amplify the *Rara* wild-type (+, 371 bp-long), the *Rara* L2 allele (427 bp-long), and the *Rara* L-(357 bp long) alleles. (H) DNA was genotyped using primers 5’-TGCTTAGCATACTTGAGAAC-3’, 5’-ACCGCACGAC ACGATAGGAC-3’ and 5’-GTAGATGCTGGGAATGGAAC-3’ to amplify the *Rarg* wild-type (+, 707 bp-long), the *Rarg* L2 (768 bp-long) and the *Rarg* L-(495 bp-long) alleles. (I) RT-qPCR analysis comparing the levels and distributions of *Sycp3*, *Rec8* and *Stra8* mRNAs in single germ cells from control and mutant ovaries at E13.5 and E14.5. The Violin plot width and length represent respectively the number of cells and the range of expression (Log2Ex). The box-and-whisker plots illustrate medians, ranges and variabilities of the collected data. The histograms show the percentages of expressing cells in each group. *Sycp3*, *Rec8* and *Stra8* are present in 94%, 91% and 72% of mutant germ cells at E14.5.

To further investigate expression of the meiotic program at the cellular level in the absence of RARs, we used RT-qPCR on single cells isolated from ovaries at E13.5 (25 control and 43 mutant germ cells) and E14.5 (40 control and 47 mutant germ cells). Consistent with IHC analyses, the proportion of germ cells expressing *Sycp3* and *Rec8*, as well as their individual levels of expression, were similar in control and mutant ovaries (Fig. 3I). In addition, the cellular expression of 12 other meiosis-specific genes including some of the ATRA-dependent class 1 to 3 genes^22^ (*Dmc1*, *Mei1*, *Meiob*, *Prdm9*, *Smc1b*, *Spo11*, *Stag3*, *Syce1*, *Syce2*, *Sycp1*, *Sycp2*, *Ugt8a*) was similar, whether RAR were present or absent (Suppl. Figs. 4–6). This confirmed the commitment of mutant germ cells toward meiosis, despite their lack of RARs.

Importantly, the meiotic gatekeeper STRA8^23^ was also expressed in germ cells of mutant ovaries, notably in those which underwent cre/ERT^2^-mediated recombination (Fig. 3C,F), albeit their number was about half-less that observed in control ovaries (Fig. 2C). Accordingly, *Stra8* mRNA was present in only 30% of the germ cells from mutant ovaries at E13.5, versus 92% in control ovaries. In addition, its expression level was slightly decreased, when compared to the control situation (Fig. 3I). This difference was smoothed out at E14.5 in mutant germ cells where the expression level of *Stra8* almost reached that measured in control germ cells at E13.5 (Fig. 3I). These observation suggested that *Stra8* expression might be delayed in the absence of RARs.

### Ablation of RARs at an earlier developmental stage dramatically impacts embryonic development but not meiosis initiation

When treated by TAM at 10.5, the mutant fetuses displayed, at E14.5, some of the eye defects typically observed in *Rarb*^−/−^;*Rarg*^−/−^ knockout fetuses (Suppl. Fig. 7). However, overall these mutant fetus were much less malformed than expected from previous works^24^. For instance, the cardiac defects described in *Rara*^−/−^;*Rarg*^−/−^ knockout fetuses^25^ were not observed. To test for the possibility that this discrepancy could be due to the timing of RARs ablation, we treated pregnant females with TAM at E9.5, and analyzed the phenotypes induced in fetuses at E14.5. The mutants generated in this way displayed most of the congenital malformations that are hallmarks of the loss of RARs^24^ (Fig. 4). We conclude therefore that deletion of RARs with TAM at E9.5 recapitulates the pathological phenotypes observed in the compound knockouts of RARs (i.e., in *Rara*^−/−^;*Rarb*^−/−^, *Rara*^−/−^;*Rarg*^−/−^ and *Rarb*^−/−^;*Rarg*^−/−^ fetuses).

**Fig. 4.**
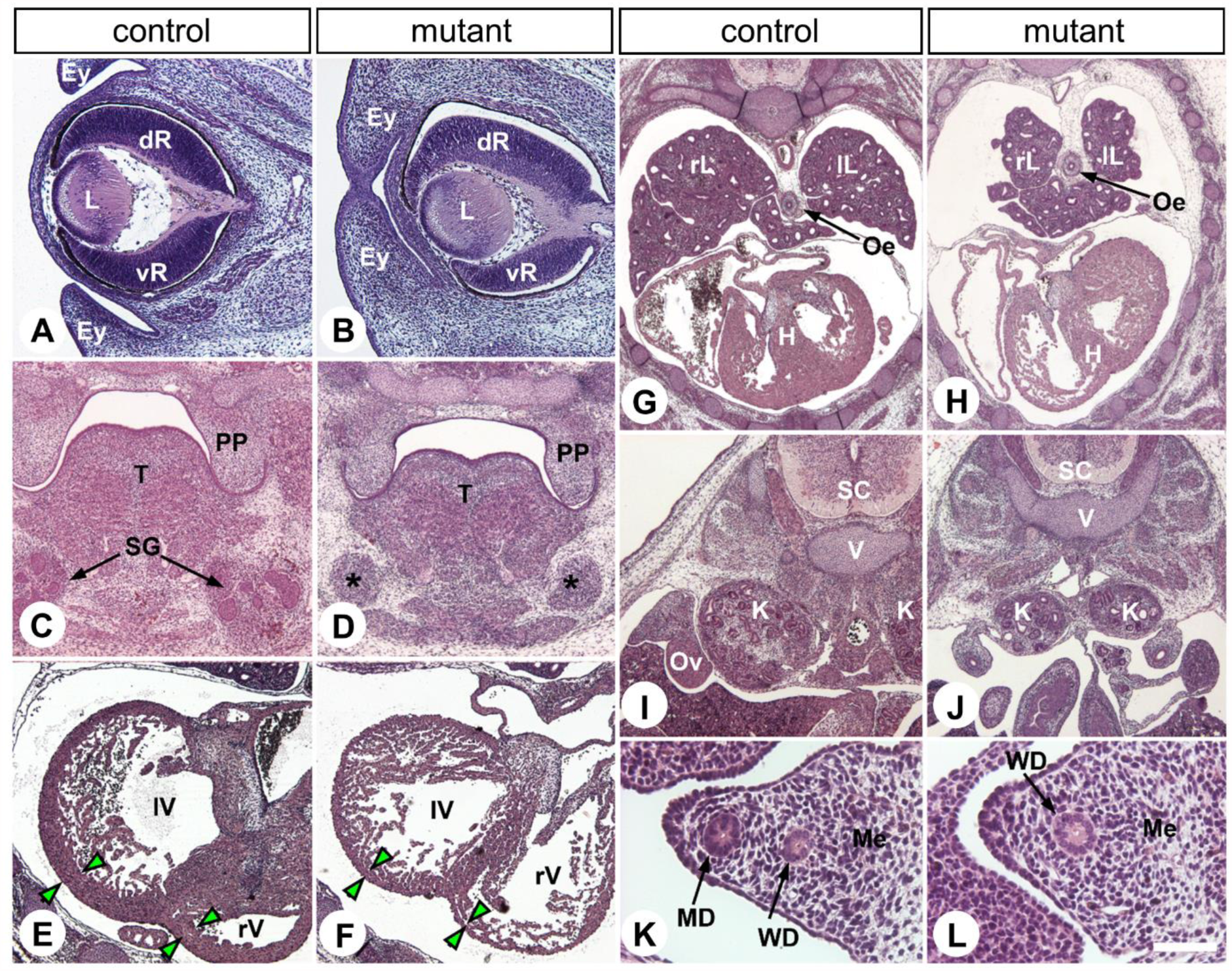
Mutant fetuses generated upon TAM injection at E9.5 display a spectrum of congenital defects typically observed at E14.5 in compound RAR-knockout mutants. Frontal histological sections at similar levels of E14.5 control (A,C,E,G,I,K) and mutant (B,D,F,H,J,L) female fetuses. (A,B) In mutants, the ventral portion of the retina (vR) is reduced in size in comparison to the dorsal retina (dR), the lens (L) is rotated ventrally and the eyelid folds (Ey) are fused together. (C,D) Mutants display mesenchymal condensations indicating the position of the salivary glands (asterisks in D), but their epithelial portion (SG in C) is absent. (E,F) In mutants, the thickness of the compact layer of the myocardium (green arrowheads) is markedly reduced in both right and left ventricles (rV and lV, respectively). (G,H) Mutants display hypoplasia of the right and left lungs (rL and lL, respectively). (I,J) Mutants, have hypoplastic kidneys (K). (K,L) Mutants lack the Müllerian duct (MD in K). Sections were stained with hematoxylin and eosin. H, heart; Me, mesonephros; Oe, oesophagus; Ov, ovary; PP, palatal process; SC, spinal cord; SG, salivary glands; T, tongue; V, vertebra; WD, Wolffian duct. Scale bar (in L): 160 µm (A,B), 320 µm (C,D,E,F,I,J), 640 µm (G,H) and 60 µm (K,L).

Given the role assigned to ATRA in sex determination^26^ and primordial germ cells proliferation^27,28^, one might have expected that deleting RARs at E9.5 would affect the gonads more dramatically than at E10.5. Yet, ovaries formed in mutants and contained normal amounts of germ cells, out of which many of them were meiotic, as indicated by their expression of SYCP3 at E14.5 (Suppl. Fig. 8).

To test for the possibility of a delay in meiosis entry in the absence of RAR (see above), we analyzed at E15.5 control and mutant ovaries of fetuses treated by TAM at E9.5. Virtually all germ cells had entered meiosis in the mutant ovaries, as attested by the detection of SYCP3 in almost all germ cells (Fig. 5A,B). Moreover, a majority of the germ cells showed thread-like segments of synaptonemal complexes positive for only SYCP3 or SYCP1 and SYCP3, indicating that they were at the late leptotene to zygotene stages (Fig. 5C,D). Both REC8 and STRA8 were detected at E15.5 in a few germ cells from control and mutant ovaries (Fig. 5E-H), as expected from previous studies^29,30^. Altogether, our results indicate that RARs are indispensable neither to enter meiosis, nor to express STRA8 in germ cells. Importantly, no increase of apoptosis was evidenced in the mutant ovaries, excluding thereby the possibility that some of the RAR-deficient germ cells experienced cell-death. We conclude that in females, RARs are not required for gonad differentiation nor for meiosis, up to the zygotene stage.

**Fig. 5.**
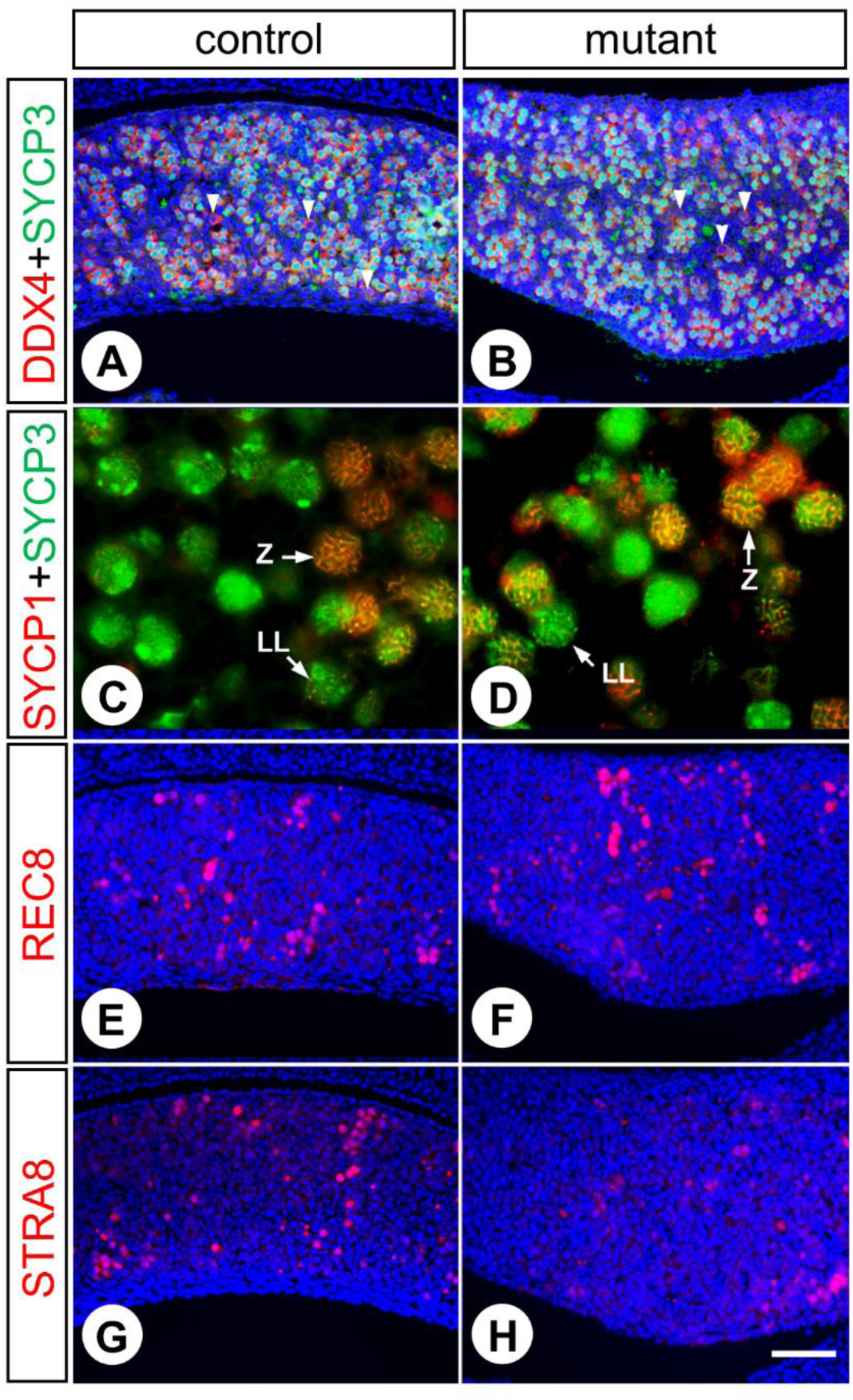
All germ cells have entered meiosis at E15.5 in ovaries of mutants lacking RARs. (A,B) Detection of germ cells (DDX4-positive, red cytoplasmic signal) expressing SYCP3 (green nuclear signal): in both control (A) and mutant (B) ovaries, almost all germ cells express SYCP3 at high levels; exceptions (i.e, cells expressing SYCP3 at low to undetectable levels) are indicated by white arrowheads. (C,D) High power magnification views of germ cells immunostained for detection of SYCP1 (red signal) and SYCP3 (green signal): thread-like structures of only SYCP3 are characteristic of late leptotene stages (LL), while those containing also SYCP1 indicate zygotene stages (Z). (E-H) Detection of REC8 (red nuclear signal) or STRA8 (red nuclear signal): both control (E,G) and mutant (F,H) ovaries contain similar small numbers of germ cells expressing REC8 or STRA8. Note that (A,E,G) and (B,F,H) are adjacent longitudinal sections through the same control and mutant ovary, respectively. Nuclei are counterstained with DAPI (blue signal). Scale bar (in H): 60 µm (A,B,E-H) and 15 µm (C,D).

## Discussion

It is widely believed that ATRA synthesized by the mesonephros is an essential paracrine factor diffusing into the ovary to trigger the differentiation of oogonia to oocytes, which progress into meiosis^13,31–36^. Nonetheless, germ cells can initiate meiosis in the absence of ALDH1A2, the ATRA-synthesizing enzyme detected in the mesonephros^7^. This seemingly contradictory data has been the source of heated and passionate debates^14,37^ but the discovery that the weak ATRA-synthesizing enzyme ALDH1A1 is capable of generating some ATRA in the developing mouse ovary^8,38^ tailored an explanation for the finding that ovarian germ cells remain able to enter meiosis in *Aldh1a2*^−/−^ knockout fetuses^8,35^. We designed the present genetic study, in which ATRA-receptors are deleted, to address by another mean the question as to whether or not ATRA is required to meiosis initiation in the mouse female germ cells.

Our results clearly show that meiotic cells expressing STRA8, REC8 and SYCP3 are present in ovaries lacking RARs. Both the number of STRA8-positive cells and the level of *Stra8* expression are lower in mutant than in control ovaries at E13.5. However, this difference diminishes at E14.5, and at E15.5 the proportion of oocytes that are at the late leptotene and zygotene stages is similar in mutant and control ovaries. It is therefore reasonable to propose that RARs participate to the timed expression of *Stra8* in the fetal ovary but, are actually not indispensable. Actually, other transcription factors such as MSX1, MSX2^39^ and DMRT1^40^, other signaling pathways such as BMPs^39,41^ and Activin A^42^, as well as the epigenetic status of chromatin^43,44^ appear more important than ATRA for proper *Stra8* expression in the fetal ovary.

More intriguingly, single cell RT-qPCR analyses at E13.5 reveal that some control and most RAR-null oocytes express meiotic genes such as *Dmc1*, *Smc1b*, *Spo11*, *Stag3*, *Syce1*, *Syce2*, before expressing *Stra8*. Nonetheless, meiosis progresses up to the zygotene stage at E15.5, despite the delay in *Stra8* expression. This finding suggests that *Stra8* expression is not be the primary event triggering meiosis in oocytes, which is unexpected given its assigned central role in this process^23^. Accordingly, *Stra8*-deficient ovarian germ cells can grow and differentiate into oocyte-like cells, without premeiotic chromosomal replication, synapsis and recombination^45^. Analysis of RAR-deficient oocytes beyond zygotene would certainly be informative, but is unfortunately not possible because RAR-null mutant fetuses die at birth from multiple congenital abnormalities^24^.

The finding that RARs are crucial for neither *Stra8* expression nor meiosis initiation implies that ATRA is not the molecule triggering meiosis, provided it does not act through a RAR-independent mechanism. Several reports describe binding of ATRA to nuclear receptors other than RARs. First, peroxisome proliferator activated receptor delta (PPARD) is activated by ATRA when the latter is delivered bound to FABP5. The interaction of ATRA with PPARD has a Kd of 15–50 nM^46^, more than two order of magnitude weaker than that of RAR, which is in the 0.1–0.2 nM range^47,48^. This concentration of ATRA is much higher than that observed in the fetal mouse ovary^7^ or required for *Stra8* expression^36^. In addition, *Fabp5* is not expressed in the mouse fetal ovary^16^, and *Ppard*-null females are fertile, indicating normal meiosis^49,50^. Second, the chicken ovalbumin upstream promoter transcription factor (NR2F2) and the testicular receptor 4 (NR2C2) can be activated by ATRA, but at non-physiological concentrations of 10-30 µM^51,52^. Third, ATRA binds to the retinoic acid receptor-related orphan receptor beta (RORB) with a Kd of 0.3 µM, but *Rorb*-null females are fertile^53^. Aside from nuclear receptors, cellular retinoic acid binding protein 1 (CRABP1) is also able to mediate some non-genomic activity of ATRA at a concentration of 100 nM^54^. However, CRABP1-null mice are fertile, and their ovaries appears fully normal^55–57^. Since all the mechanisms identified to date through which ATRA could act independently of RARs can be excluded, ATRA is not the meiosis inducing substance. In agreement with this view, the report by Chassot et al. (accompanying manuscript) shows that simultaneous ablation of *Aldh1a1*, *Aldh1a2* and *Aldh1a3* in either the somatic cells or in the whole ovary of mice do not impair *Stra8* expression and meiosis initiation. These findings are in keeping with the recent observation that initiation of meiosis and expression of *Stra8* by spermatocytes occurs without ATRA^58^.

Our results apparently contradict previous observations^13,31–36^, but several explanations can be proposed to reconcile the discrepancies. First, experiments performed using BMS-204493^6,29,41^ and AGN193109^5,8,26,38^ to impair ATRA signaling in fetal ovaries or testes must be interpreted with caution. Actually these ligands are not RAR antagonists, as they are usually referred to as, but pan-RAR inverse agonists, which are capable of repressing RAR basal activity by favoring and stabilizing recruitment of corepressors, even in the absence of an endogenous agonistic ligand such as ATRA^59–62^. As RAR binding sites are present in the *Stra8* promoter on which RARs are actually bound *in vivo*^7,63,64^, adding a pan-RAR inverse agonist predictably induces recruitment of NCOR1 and NCOR2 corepressors at the locus, promotes chromatin compaction^65^ and thereby *Stra8* extinction, irrespective of the presence of ATRA in the ovary. The same holds true for the *Rec8* gene, which also contains a RARE^63,66^. The fact that expression of these two genes is artificially shut down upon exposure to a pan-RAR inverse agonist does not mean that their expression is normally controlled by endogenous ATRA. If one assumes that *Rec8* and *Stra8* are actually not regulated by endogenous ATRA, it is perfectly logical to find them expressed in germ cells lacking RARs (present study). For the same reason, the fact that AGN193109 abrogates the ectopic expression of *Stra8* in *Cyp26b1-*null mouse testes is not a proof that expression of *Stra8* depends on ATRA signaling. Therefore it cannot be used as an argument to quash the theory proposed by Kumar et al. in 2011 that a molecule distinct from ATRA is involved in the initiation of meiosis^8^. Second, it has been recurrently emphasized that alterations generated by exogenously administered ATRA do not necessarily reflect physiological processes^24,67–69^. In the case of the gonads, the simple fact that RAR-binding sites are present in *Stra8* and *Rec8* genes can explain their forced expression and initiation of meiosis by supra-physiological concentrations of ATRA added to fetal rat ovaries^70^ and to mouse testes cultured *in vitro*^5,6^.

Involvement of ATRA in meiosis is commonly interpreted according to the prevailing idea according which both a MIS, ATRA, and a MPS, the ATRA-degrading enzyme CYP26B1, are required to account for the sex-specific timing of meiotic initiation^13,14^. Since the present work disqualifies ATRA as an MIS, the “MPS-only” hypothesis^71^ would reconcile the discrepancies present in the literature: germ cells are programmed to initiate meiosis unless prevented from doing so by an MPS produced in the testis^72^. According to this scenario, inducing mitotic quiescence instead of meiosis in male germ cells during fetal life requires a yet unknown substance, the MPS, which is undoubtedly produced by CYP26B1^7,73^. Efforts should now be put to identify this substance.

## Methods

### Mice and treatments

Mice were on a mixed C57BL/6 (50%)/129/SvPass (50%) genetic background. They were housed in a licensed animal facility (agreement #C6721837). All experiments were approved by the local ethical committee (Com’Eth, accreditations APAFIS#5638-2016061019045714 and APAFIS#5639-201606101910981), and were supervised by N.B.G., M.M. and N.V., who are qualified in compliance with the European Community guidelines for laboratory animal care and use (2010/63/UE). To inactivate *Rar* coding genes, mice bearing *lox*P-flanked (L2) alleles of *Rara*^74^ and *Rarg*^75^ and null (L-) alleles of *Rarb*^76,77^ were crossed with mice bearing the ubiquitously expressed, tamoxifen-inducible, cre/ERT^2^ recombinase-coding *Tg*(*Ubc-cre/ERT*^2^ transgene^*78*^. Females homozygous for L2 alleles of *Rara* and *Rarg* and for L-alleles of *Rarb* (i.e., *Rara*^L2/L2^;*Rarg*^L2/L2^;*Rarb*^L-/L−^) were mated with males bearing one copy of the *Tg*(*Ubc-cre/ERT*^2^), and homozygous for L2 alleles of *Rara* and *Rarg* and for L-alleles of *Rarb* (i.e., *Tg*(*Ubc-cre/ERT^2^*);*Rara*^L2/L2^;*Rarg*^L2/L2^;*Rarb*^L-/L−^). Noon of the day of a vaginal plug was taken as 0.5 day embryonic development (E0.5). To activate the cre/ERT2 recombinase in embryos, one tamoxifen (TAM) treatment (130 mg/kg body weight) was administered to the pregnant females by oral gavage at E9.5 or at E10.5. TAM (T5648, Sigma-Aldrich) was dissolved in ethanol at a concentration of 100mg/mL and further diluted in sunflower oil to a concentration of 10mg/ml. This resulted in embryos or fetuses null for *Rarb* in which *Rara* and *Rarg* were ablated upon TAM induction when they were bearing *Tg*(*Ubc-cre/ERT^2^*) (referred to as mutants), as well as their control littermates when the embryos or fetuses were free of *Tg*(*Ubc-cre/ERT^2^*) (referred to as controls). Importantly, embryos, fetuses and females null for *Rarb* display normal ovaries and are totally fertile^76,77^. To assess for cre/ERT^2^-directed excision, we introduced the *Gt(ROSA26)^ACTB-tdTomato-EGFP^* reporter transgene (referred to as *mT/mG*), which directs expression of a membrane-targeted green fluorescent protein (mGFP) in cells that have experienced cre-mediated deletion^79^, in the *Tg*(*Ubc-cre/ERT*^2^);*Rara*^L2/L2^;*Rarg*^L2/L2^;*Rarb*^L-/L−^ genetic background. Embryos and fetuses were collected by caesarean section and the yolk sacs, tail biopsies or female gonads were taken for DNA extraction. Genotypes were determined as described^74–76,78,79^.

### External morphology, histology and immunohistochemistry

Following collection, E11.5 embryos to E15.5 fetuses were fixed overnight in cold 4% (w/v) paraformaldehyde (PFA) in phosphate buffered saline (PBS). After removal of the fixative, embryos and fetuses were rapidly rinsed in PBS and placed in 70% (v/v) ethanol for long-term storage and external morphology evaluation. They were next embedded in paraffin. Consecutive, frontal, 5 µm-thick sections were made throughout the entire specimens. For histology, sections were stained with hematoxylin and eosin (H&E). For immunohistochemistry (lHC) antigen were retrieved for 1 hour at 95°C either in 10 mM sodium citrate buffer at pH 6.0 or, only in the case of IHC for detection of REC8 and SYCP1, in Tris-EDTA at pH 9.0 [10 mM Tris Base, 1 mM EDTA, 0.05% (v/v) Tween 20]. Sections were rinsed in PBS, then incubated with appropriate dilutions of the primary antibodies or a mixture of them (i.e., anti-DDX4 and anti-SYCP3; anti-SYCP1 and anti-SYCP3) in PBS containing 0.1% (v/v) Tween 20 (PBST) for 16 h at 4°C in a humidified chamber. After rinsing in PBST (3 times for 3 min each), detection of the bound primary antibodies was achieved for 45 min at 20°C in a humidified chamber using Cy3-conjugated donkey anti-rabbit, Alexa Fluor 488-conjugated donkey anti-mouse or goat anti-chicken antibodies, depending on the origin of the primary antibody depending on the origin of the primary antibody (Suppl. Table 1). Nuclei were counterstained with 4’,6-diamidino-2-phenyl-indole (DAPI) diluted at 10 µg/ml in the mounting medium (Vectashield; Vector).

### Characterization and counts of the meiotic germ cells

Germ cell counts were performed on pairs of fetuses consisting of one mutant and one of its control littermates, and each experiment was repeated on at least 3 different fetuses per genotype, age and stage of TAM treatment. The total number of germ cells in ovaries of control and mutant E14.5 fetuses was quantified using immunostaining for DDX4. Meiotic germ cells were quantified using immunostaining for SYCP3, REC8 and STRA8. A double-immunostaining for SYCP1 and SYCP3 was performed to detect germ cells at the zygotene stage. Data were expressed as percentages related to the number of DDX4-positive cells. Statistical analysis was done by a two-tail Student *t*-test, assuming unequal variances after arcsine transformation of the percentages of germ cells expressing the meiotic markers.

### Single cell RT-qPCR and data processing

Gonads from E13.5 and E14.5 control and mutant fetuses were dissected out in PBS and sexed by their appearance under the microscope. Ovaries were separated from mesonephros using the cutting edge of a 25G needle. Dissociated cells were obtained by incubating the gonads for 15 minutes in PBS containing 0.5 µM EDTA^80^. To allow cell pricking, gonads were then transferred in PBS containing 4 mg/ml BSA. Cells were released by puncture of the gonad with 25G needles. This technique generally resulted in a suspension of germ cells, which are recognized as large, refringent, cells, with low somatic cell contamination^81^. Cells (n=88 at E13.5 and n=132 at E14.5), were collected individually using pulled Pasteur pipets, and transferred as isolated, single, cells into micro-tubes containing 5 µl of cold 1X SuperScript IV VILO Master Mix for two-step RT-qPCR containing 6 units RNasin (Promega) and 0.5% (v/v) NP40. The micro-tubes were then stored at −80°C for later processing. Two-step, single-cell, gene expression analysis was achieved on the BioMark HD system (Fluidigm) using SsoFast EvaGreen Supermix with low ROX (Bio-Rad Laboratories) according to manufacturer’s instructions. The set of primers used for qPCR are listed (Suppl. Table 2). Cts were recovered and analyzed by the Fluidigm Real-Time PCR Analysis software, using the linear (derivative) baseline correction method and the auto (global) Ct threshold method. Limit of detection (LOD) was set to a Ct of 28. The quality threshold was set to 0.65. Cts for qPCR reactions that failed the quality threshold (melting curves with deviating Tm temperatures) were converted to 28. The Ct values were then exported to Excel and converted to expression levels using the equation Log Ex = Ct[LOD]-Ct[Assay]. The expressions of *Actb* and *Tbp* housekeeping genes were measured to determine which sample actually contained a cell. Cells that fail to amplify systematically all the tested genes or to express germ cell markers (*Dazl*, *Ddx4* and *Kit*) were excluded from further analysis. The percentages of cells excluded was 23% at E13.5 and 34% at E14.5. This can be explained by the greater difficulty to isolate and collect germ cells at E14.5, compared to E13.5. The final analysis was done using n=68 individual cells at E13.5 (n=25 controls, n=43 mutants) and n=87 individual cells at E14.5 (n=40 controls, n=47 mutants). Results were not normalized relative to housekeeping genes, as their expression did not differ between control and mutant cells. Percentages of cells expressing a gene of interest are represented as histograms, and expression levels are represented as violin and box plot using R Studio software.

## Acknowledgments

We thank Dr Amandine Chassot, Dr Marie-Christine Chaboissier and Dr Eric Pailhoux for discussions, advices and critical reading of the manuscript. We also thank Dr Christelle Thibault-Carpentier from the Genomeast platform for her input (http://genomeast.igbmc.fr/). This work was supported by grants from CNRS, INSERM, UNISTRA and Agence Nationale pour la Recherche (ANR; 10-BLAN-1239, 13-BSV6-0003 and 13-BSV2-0017), as well as from EU (FP7-PEOPLE-IEF-2012-331687). Their studies were also supported in part by the grant ANR-10-LABX-0030-INRT, a French State fund managed by the ANR under the frame programme Investissements d’Avenir labelled ANR-10-IDEX-0002-02.

## Author contributions

N.V., M.M., and N.B.G. designed the study, analyzed the data and wrote the paper. N.V., M.M., D.C., B.F., M.K., V.A. and M.T. performed the experiments, analyzed the data, and discussed the results. M.T. additionally commented on the manuscript.

## Additional information

**Supplementary Information** accompanies his paper.

## Supplementary Information

**Suppl. Fig. 1.**
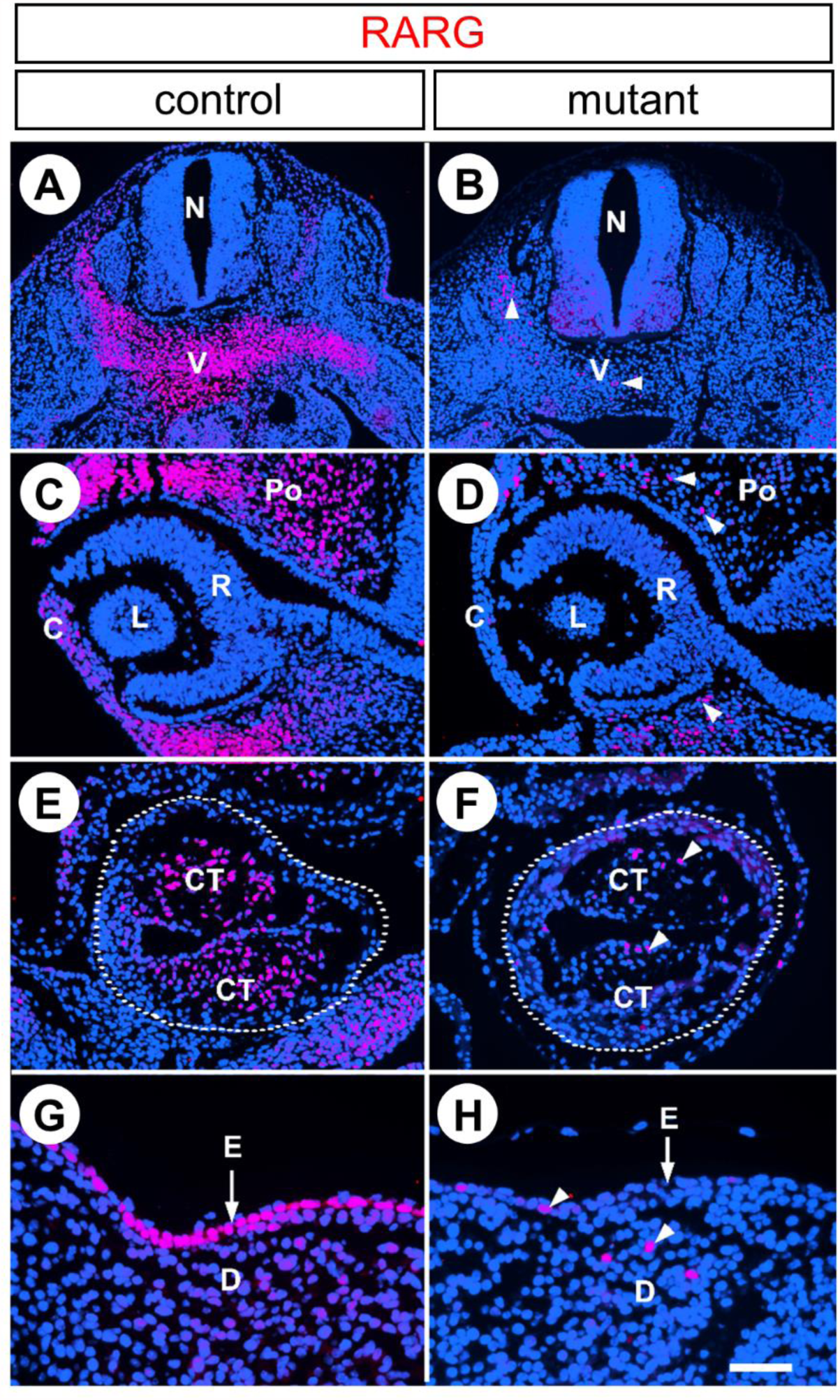
Excision of RARG after administration of TAM at E10.5. immunohistochemical detection of RARG (red signal) on frontal histological sections at similar levels of control and mutant embryos at E11.5, namely 24 hours after TAM administration. (A,C,E,G) RARG is strongly expressed in precartilaginous vertebrae, in periocular and corneal mesenchyme, in conotruncal ridges and in epidermis of the control embryo. (B,D,F,H) Expression of RARG is nearly abolished in the tissues of the mutant embryo. Nuclei are counterstained with DAPI (blue signal). C, cornea; CT, conotruncal ridges; D, dermis; E, epidermis; L, lens; Po, periocular mesenchyme; R, retina; V vertebra; The dotted lines mark off the periphery of the heart outflow tract. White arrowheads indicate nuclei that are still expressing RARG in the mutant tissues. Scale bar (in H): 160 µm (A,B), 80 µm (C-F) and 40 µm (G-H).

**Suppl. Fig. 2.**
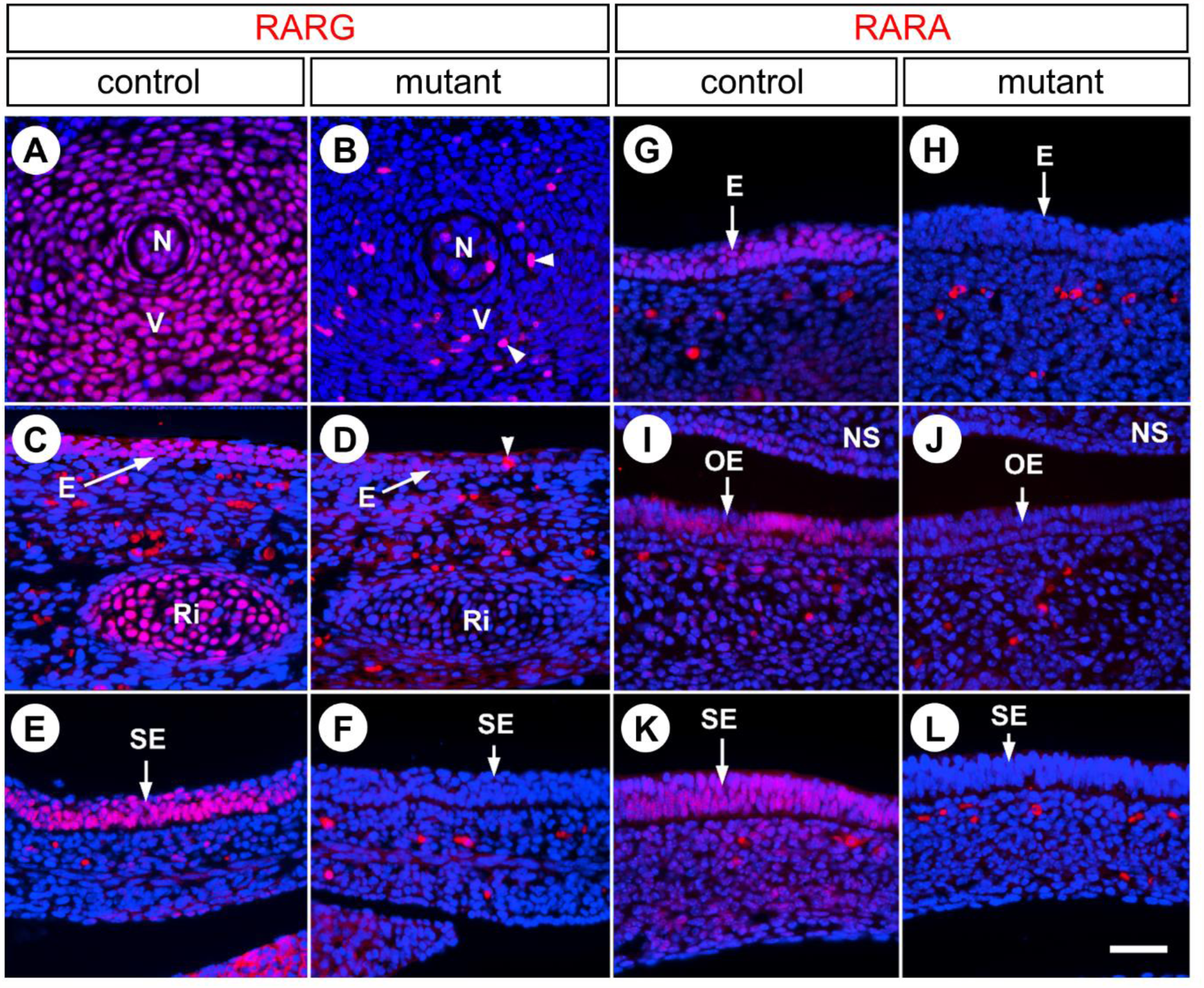
Excision of RARG and RARA after administration of TAM at E10.5. Immunohistochemical detection of RARG (red signal in A-F) and RARA (red signal in G-L) on histological sections of control and mutant fetuses at E14.5, namely 4 days after TAM administration. (A,C,E) RARG is strongly expressed in all cartilages including vertebrae (V) and ribs (Ri), as well as in the epidermis (E) and epithelium of the stomach (SE) of control fetuses. (B,D,F) Expression of RARG is abolished in almost all cells in tissues of mutant fetuses. (G,I,K) RARA is strongly expressed in the snout epidermis (E), olfactory epithelium (OE) and epithelium of the stomach (SE) of control fetuses. (H,J,L) Expression of RARA is lost in tissues of the mutant fetuses. Nuclei are counterstained with DAPI (blue signal). N, notochord; NS, nasal septum. White arrowheads indicate nuclei that are still expressing RARG in mutant tissues. Scale bar (in L): 40 µm (A-L).

**Suppl. Fig. 3.**
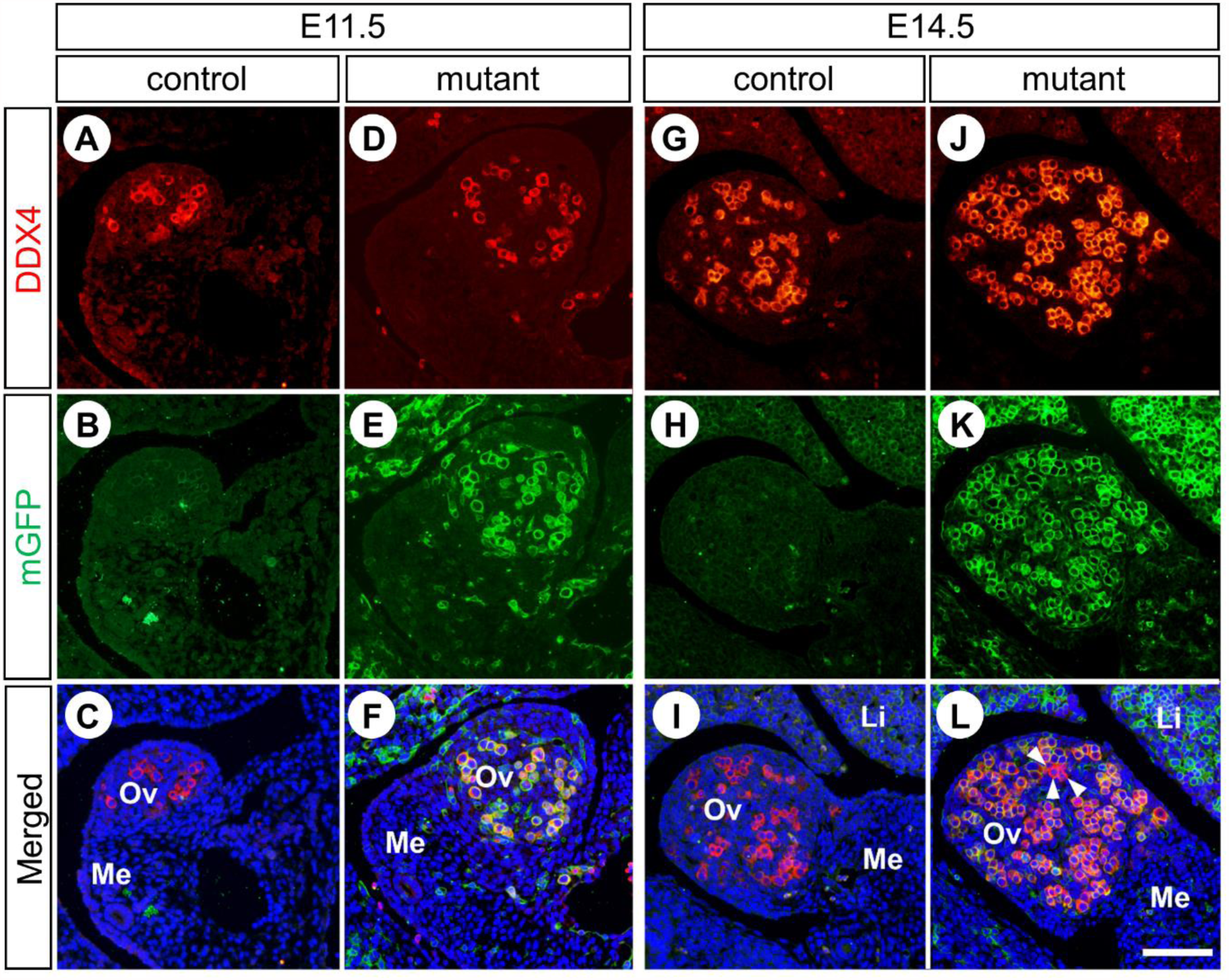
Excision of the *mT/mG* reporter in fetal ovaries after administration of TAM at E10.5. Detection of DDX4 (red cytoplasmic signal) and mGFP (green membranous signal) on histological sections of control and mutant ovaries at E11.5 (A-F) and E14.5 (G-L), namely 24 hours and 4 days after TAM administration, respectively. Efficient *mT/mG* excision is assessed by GFP expression in almost all germ cells (D-F and J-L); exceptions (i.e, red cytoplasm without a green contour) are indicated by white arrowheads. Most importantly, GFP is never detected in control ovaries, i.e., in the absence of cre/ERT^2^ (A-C and G-I). Nuclei are counterstained with DAPI (blue signal). Li, Liver; Me, mesonephros; Ov, ovary. Scale bar (in L): 80 µm (A-L).

**Suppl. Fig. 4.**
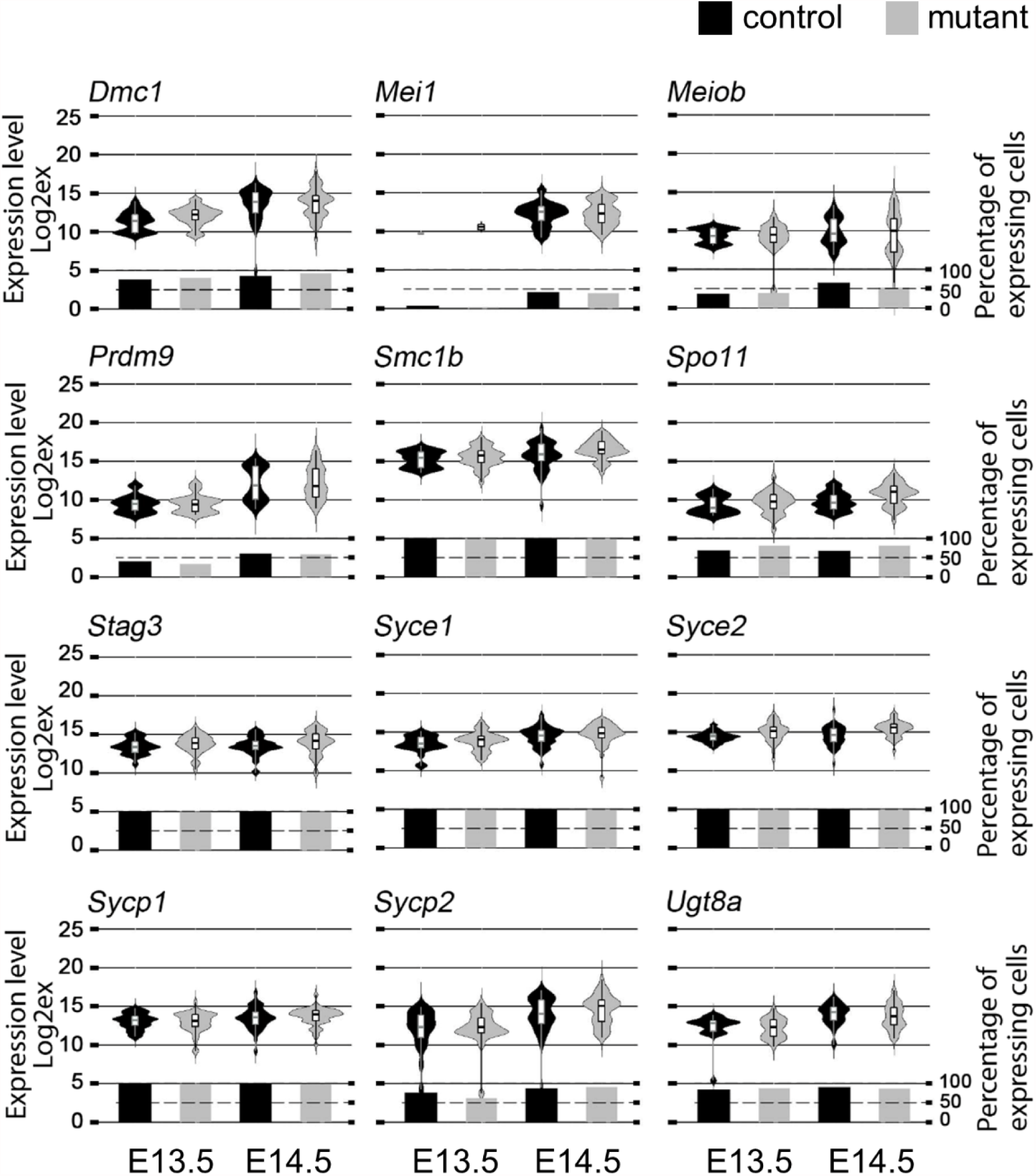
Meiotic genes are expressed at E13.5 and E14.5 in ovaries of mutant lacking RARs. RT-qPCR analysis comparing the levels and distributions of the meiotic markers *Dmc1*, *Mei1*, *Meiob*, *Prdm9*, *Smc1b*, *Spo11*, *Stag3*, *Syce1*, *Syce2*, *Sycp1*, *Sycp2* and *Ugt8a* mRNAs in single germ cells from control and mutant ovaries at E13.5 and E14.5. The Violin plot width and length represent respectively the number of cells and the range of expression (Log2Ex). The box-and-whisker plots illustrate medians, ranges and variabilities of the collected data. The histograms show the percentages of expressing cells in each group. A total of 155 cells were used for analysis (for see details, see Material and Methods).

**Suppl. Fig. 5.**
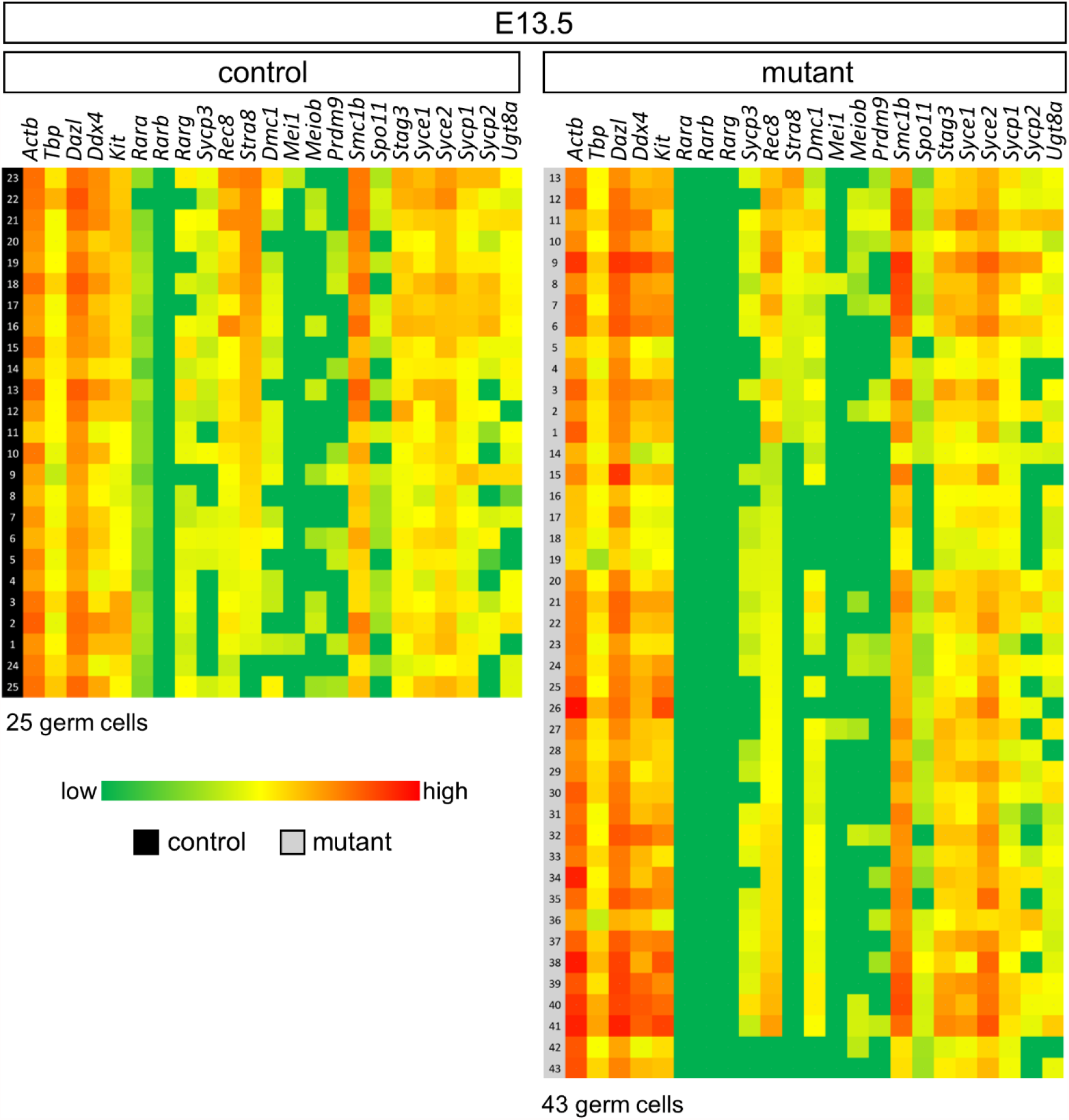
Meiotic genes are expressed at E13.5 in ovaries of mutant lacking RARs. Heatmap showing the expression of 23 genes (columns) for each of the germ cell analyzed (rows). Green, yellow and red indicate low, intermediate and high expression levels, respectively. Cell clustering was based on the *Stra8* gene expression pattern. Control cells are numbered 1 to 25 (black boxes), and mutant cells are numbered 1 to 43 (grey boxes).

**Suppl. Fig. 6.**
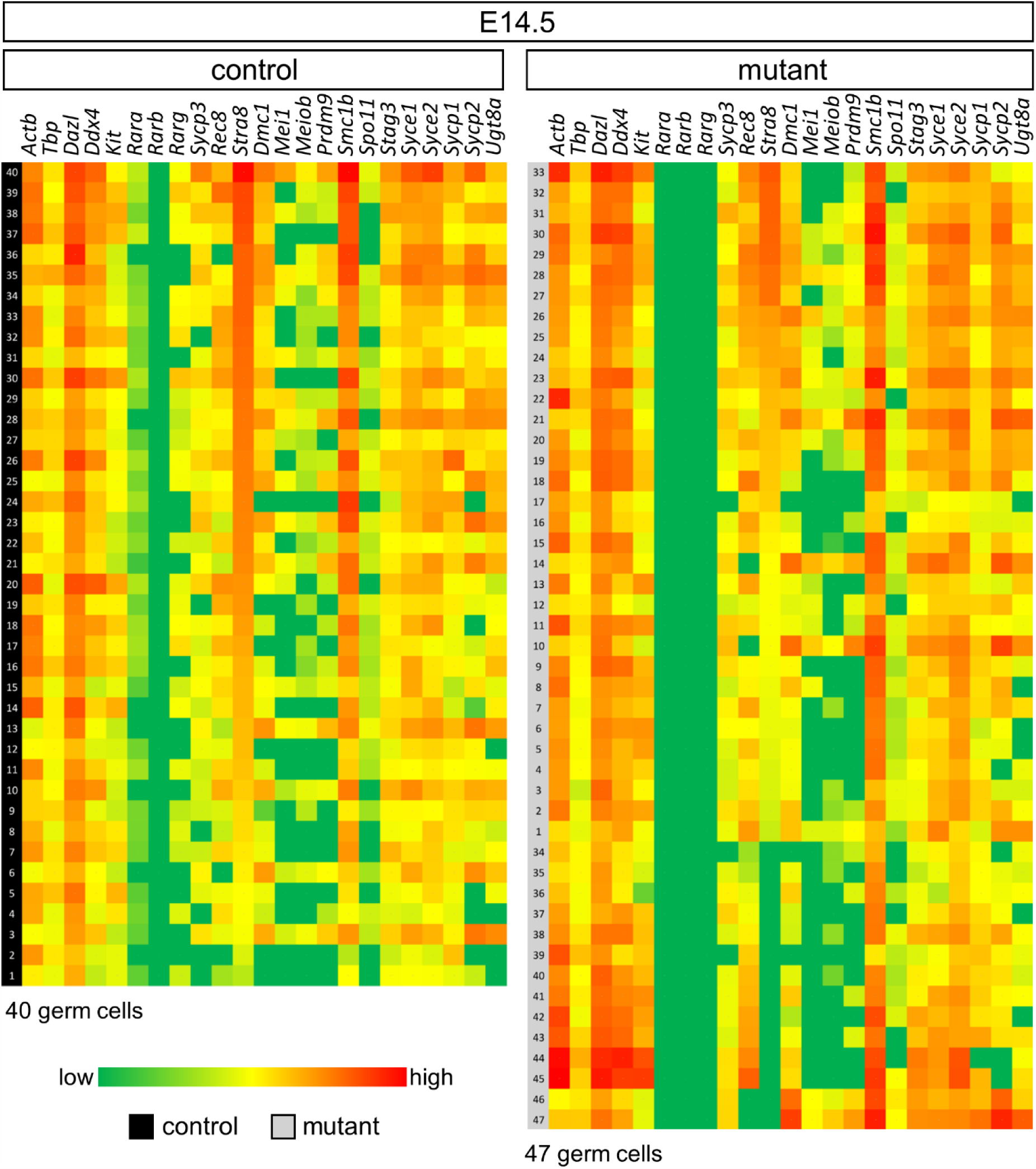
Meiotic genes are expressed at E14.5 in ovaries of mutant lacking RARs. Heatmap showing the expression of 23 genes (columns) for each of the germ cell analyzed (rows). Green, yellow and red indicate low, intermediate and high expression levels, respectively. Cell clustering was based on the Stra8 gene expression pattern. Control cells are numbered 1 to 40 (black boxes), and mutant cells are numbered 1 to 47 (grey boxes).

**Suppl. Fig. 7.**
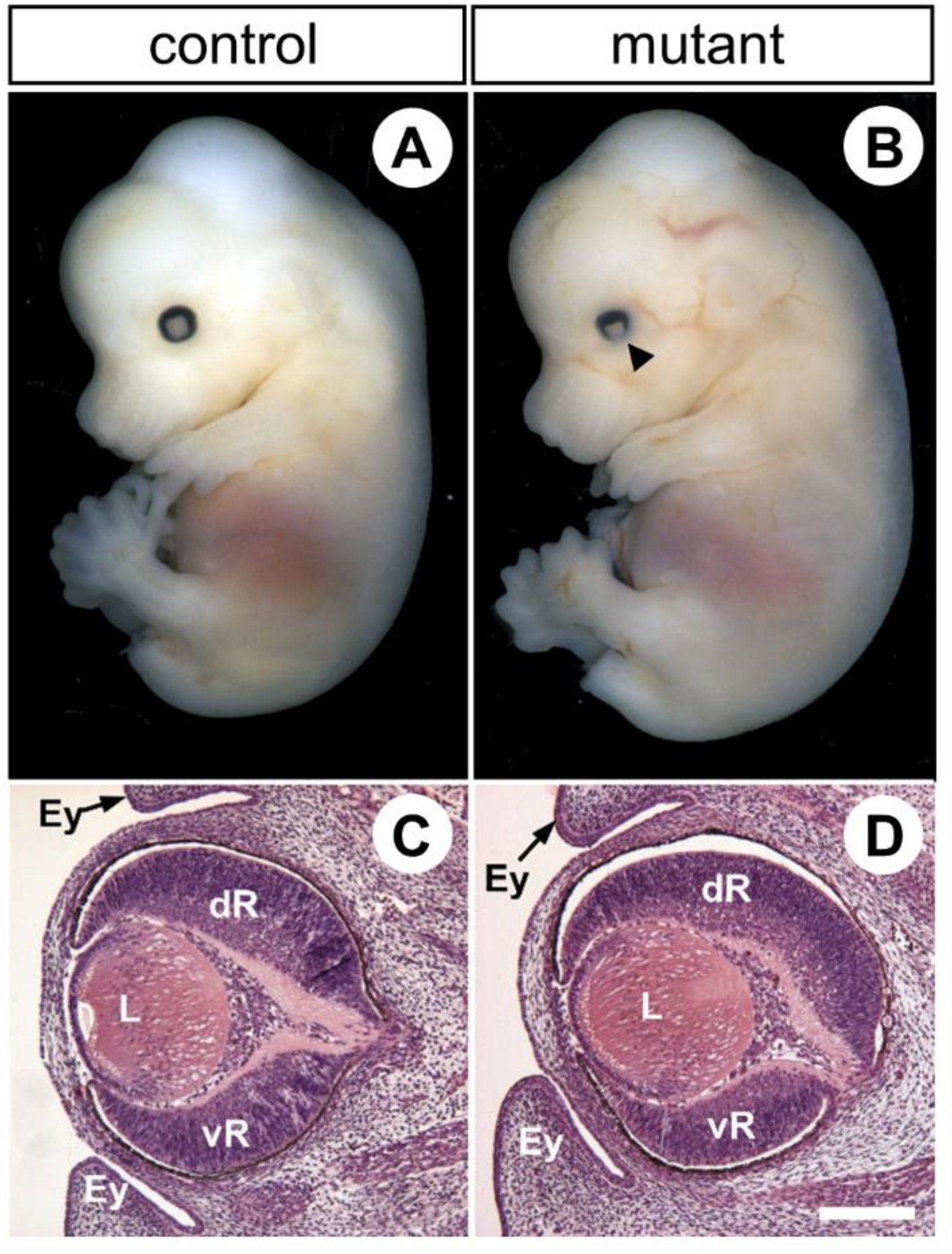
Congenital malformations in E14.5 mutant fetuses generated upon TAM injection at E10.5 are restricted to eye defects typically observed in RAR-knockout mutants. (A,B) The pigmented retina is externally less visible in the mutant (arrowhead in B). This is because the insertions of the upper and lower eyelid folds (Ey) are abnormally close to each other. (C,D) In the mutant, in addition to the closer eyelid folds, the ventral portion of the retina (vR) is reduced in size in comparison to the dorsal retina (dR) and the lens (L) is rotated ventrally. Histological sections were stained with hematoxylin and eosin. Scale bar (in D): 160 µm (C,D).

**Suppl. Fig. 8.**
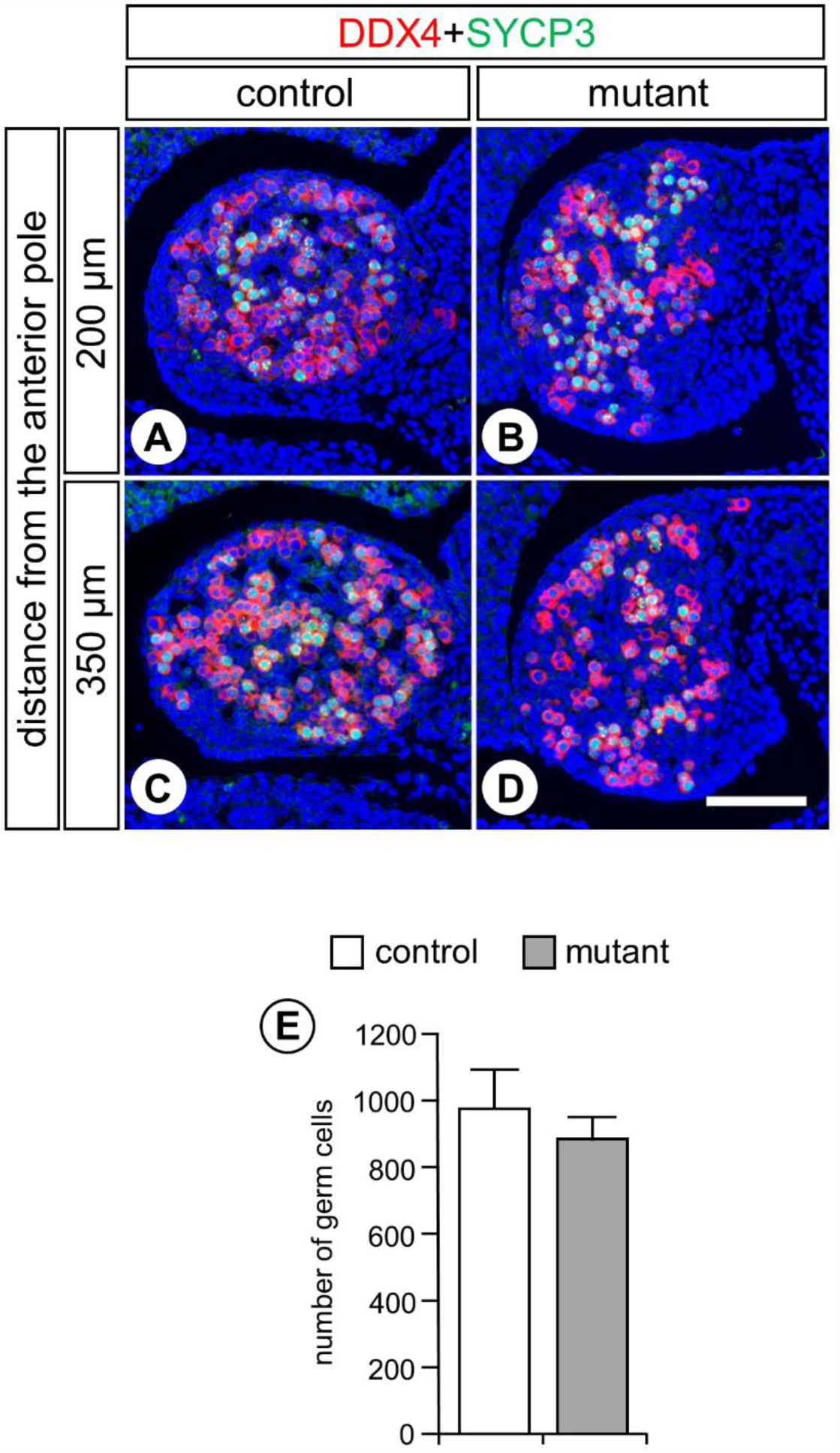
Ovaries of mutants generated upon TAM injection at E9.5 display a normal number of germ cells and contain meiotic cells. (A-D) Detection of DDX4 (red cytoplasmic signal) and SYCP3 (green nuclear signal), on transverse histological sections at two different levels of the ovaries from control and mutant fetuses at E14.5. (A) and (C), and (B) and (D), are consecutive histological sections. The positions of histological sections along the anteroposterior axis is indicated on the left in terms of distance from the anterior pole of the ovary (i.e., 200 and 350 microns). Nuclei are counterstained with DAPI (blue signal). Scale bar (in D): 60 µm. (E) Average of the total number of germ cells present at the 7 different levels of the ovaries, 75 microns apart, in 3 control (white bar) and 3 mutant fetuses (grey bar).

**Suppl. Table 1.**
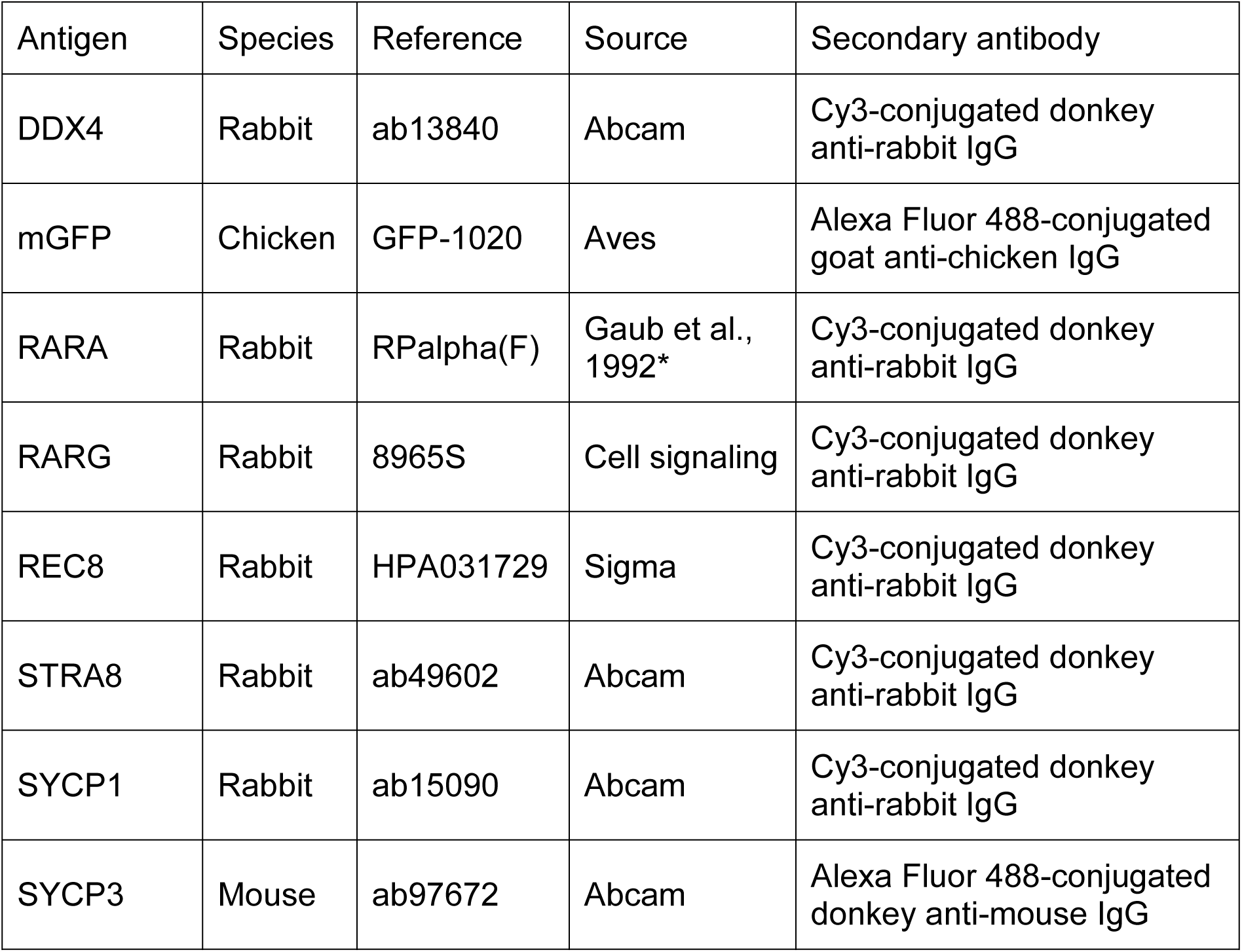
Antibodies used in immunohistochemistry experiments. *Gaub *et al.*, (1992) *Exp. Cell Res.* 201:335-346.

**Suppl. Table 2.**
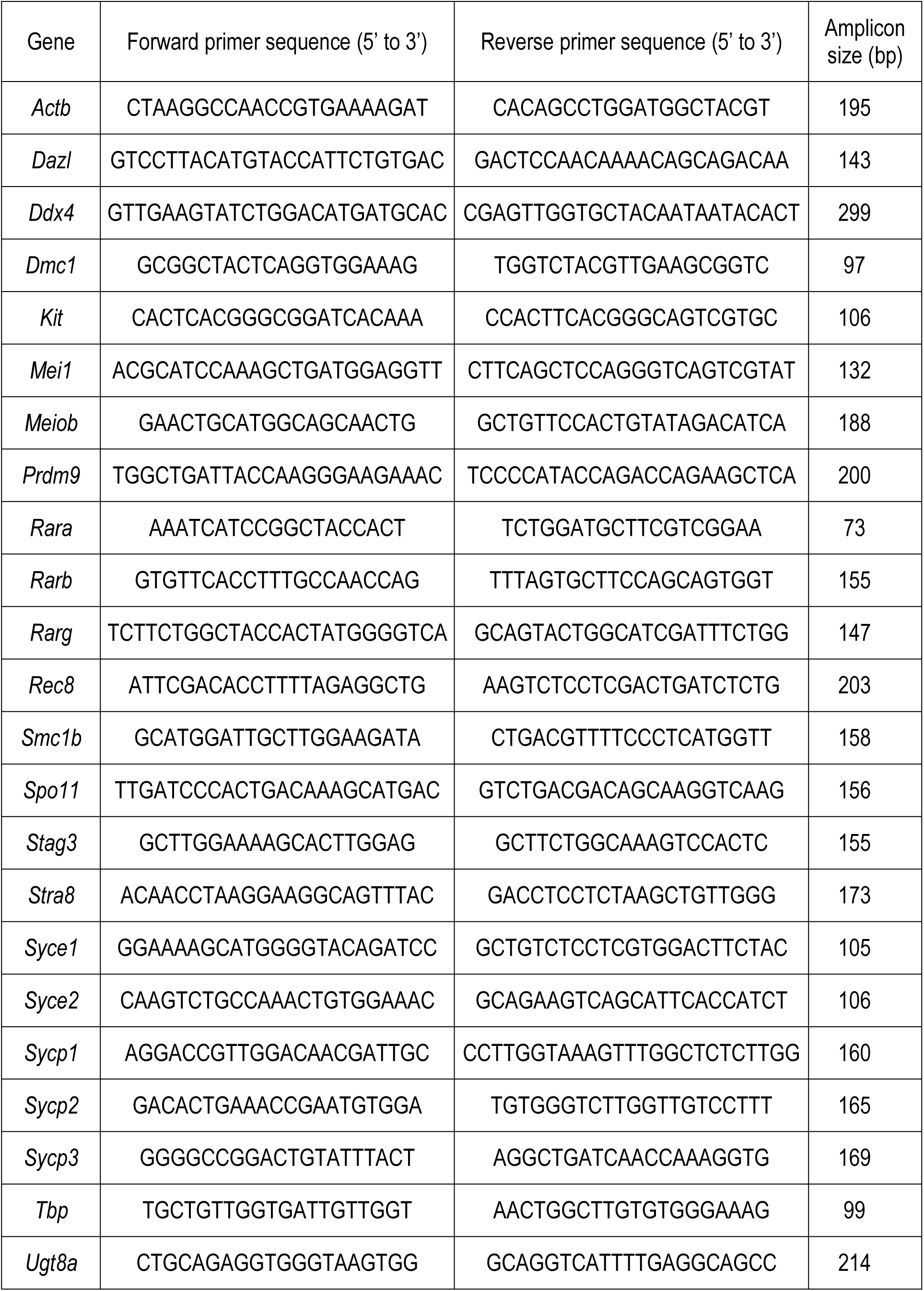
Primers used for RT-qPCR on single cells.

